# Environments with a high probability of incompatible crosses do not select for mate avoidance in spider mites

**DOI:** 10.1101/395301

**Authors:** Leonor R Rodrigues, Flore Zélé, Inês Santos, Sara Magalhães

## Abstract

Arthropods are often infected with *Wolbachia* inducing cytoplasmic incompatibility (CI), whereby crosses between uninfected females and infected males yield unviable fertilized offspring. Although uninfected females benefit from avoiding mating with *Wolbachia*-infected males, this behaviour is not present in all host species. Here we measured the prevalence of this behaviour across populations of the spider mite *Tetranychus urticae*. Females from five populations originally fully infected with *Wolbachia* showed no preference, possibly because they did not face the choice between compatible and incompatible mates in their environment. Hence, to determine whether this behaviour could be selected in populations with intermediate *Wolbachia* infection frequency, we performed 15 generations of experimental evolution of spider-mite populations under i) full *Wolbachia* infection, ii) no infection, or iii) mixed infection. In the latter selection regime, where uninfected females were exposed to infected and uninfected males at every generation, mating duration increased relative to the uninfected regime, suggesting the presence of genetic variation for mating traits. However, mate choice did not evolve. Together, these results show that CI-inducing *Wolbachia* alone does not necessarily lead to the evolution of pre-copulatory strategies in uninfected hosts, even at intermediate infection frequency.

## Introduction

Organisms are often exposed to parasites, risking severe fitness costs upon infection. Hosts are thus expected to be under strong selection to avoid being parasitized (Parker et al. 2011; Sarabian et al. 2018). This may be possible via hiding, fleeing from parasites, avoiding infected conspecifics, and/or avoiding food and habitats where encounters with parasites are likely (Schmid-Hempel 2011; Sarabian et al. 2018). Moreover, hosts may avoid mating with parasitized conspecifics. This avoidance of infection via mate choice is widespread across different host species (reviewed in Beltran-Bech and Richard 2014), and forms the basis of the Hamilton-Zuk hypothesis, which proposes that mate choice can be based upon traits associated with resistance to parasites (Hamilton and Zuk 1982).

*Wolbachia* are widespread endosymbiotic bacteria commonly found in arthropods, whose key feature is the capacity to manipulate the cellular and reproductive processes of its host, frequently leading to a decrease in host fitness (Werren et al. 2008). The most common *Wolbachia*-induced phenotype is cytoplasmic incompatibility (CI), a mechanism that results in the embryonic death of fertilized offspring from crosses between *Wolbachia*-uninfected females and *Wolbachia*-infected males. As all other crosses are compatible, CI promotes *Wolbachia* spread by indirectly (i.e. via infected males) increasing the success of infected females relative to that of uninfected females. As the number of offspring resulting from incompatible crosses is reduced relative to those of compatible ones, *Wolbachia* reduces drastically the fitness of both uninfected females and infected males. Such an adverse effect of CI is expected to exert a strong selective pressure on hosts to evolve strategies that reduce the frequency and/or costs of such matings (Charlat et al. 2003; Champion de Crespigny et al. 2005, 2006, Champion de Crespigny and Wedell 2006, 2007; Telschow et al. 2007; Sahoo 2016).

Discrimination of compatible mates prior to mating has been proposed as a potential strategy to avoid CI (Hoffmann et al. 1990; Vala et al. 2004; Champion de Crespigny and Wedell 2007). This, in turn may lead to reproductive isolation between *Wolbachia*-infected and -uninfected lineages. Indeed, different theoretical models predict that both bi-(Telschow et al. 2005) and unidirectional CI (Telschow et al. 2007) can select for premating isolation. *Wolbachia* may thus severely reduce gene flow between populations, both by decreasing the viability of crosses between infected and uninfected individuals and by selecting for mate discrimination in uninfected females or infected males (Jaenike et al. 2006; Buellesbach et al. 2014).

Several studies tested mate discrimination in species infected by CI-inducing *Wolbachia*, with variable outcomes. Indeed, whereas some studies did not find evidence for mate choice (Hoffmann and Turelli 1988; Hoffmann et al. 1990; Wade and Chang 1995; Champion de Crespigny and Wedell 2007; Duron et al. 2011; Arbuthnott et al. 2016), others found that individuals discriminate between *Wolbachia*-infected and -uninfected mates (Vala et al. 2004; Jaenike et al. 2006; Koukou et al. 2006). These contrasting results suggest that discrimination has not been universally selected across species and may also vary between populations within species. For instance, both *Wolbachia*-uninfected females and *Wolbachia*-infected males did not exhibit preference for infected or uninfected mates in a population of *D. melanogaster* and in another of *D. simulans* (Champion de Crespigny and Wedell 2007). However, a few years later, a different study in *D. melanogaster* has shown that the existence of assortative mating depends on the interaction between *Wolbachia* infection status and the genotype of the host (Markov et al. 2009).

This variation in the ability to discriminate between *Wolbachia*-infected and - uninfected individuals may hinge upon the benefits that such behaviour provides. Indeed, mate preference is expected to be selected only if individuals evolve in environments with intermediate infection frequencies, as it is only under these circumstances that incompatible matings occur and choice is possible. However, the spread *Wolbachia* might be very rapid, which limits the range of conditions under which CI can select for mate preference (Engelstädter and Telschow 2009). Hence, models predict that discrimination is more likely to evolve in populations in which *Wolbachia* induce incomplete CI, fecundity costs, and/or is imperfectly transmitted, as this slows down its spread (Champion de Crespigny et al. 2005). This should also be the case in structured host populations with migration below a critical rate, as this increases the likelihood of a stable infection polymorphism (Flor et al. 2007; Telschow et al. 2007; Engelstädter and Telschow 2009). However, although the prevalence of *Wolbachia* and the intensity of CI vary across species and populations (Gotoh et al. 2007; Hughes et al. 2011; Hamm et al. 2014; Ahmed et al. 2015; Zélé et al. 2018a), no experimental study so far has specifically controlled for the recent infection history of host populations. Thus, it is as yet unclear whether the frequency of *Wolbachia* in a population will affect the evolution of female choice towards uninfected vs infected males, as predicted by mathematical simulations (Champion de Crespigny et al. 2005; Telschow et al. 2007).

In this study, we investigate whether females from populations of the spider mite *Tetranychus urticae* vary in their choice towards males that are either infected or uninfected with *Wolbachia*. Moreover, we test whether this trait responds to selection. An earlier study found that *Wolbachia*-uninfected *T. urticae* females preferred to mate with *Wolbachia*-uninfected males and that they preferentially oviposit near uninfected eggs, increasing the chances that their offspring would engage in compatible matings (Vala et al. 2004). However, this study was done with a single isogenic line. We thus tested the generality of this finding by studying whether *Wolbachia*-uninfected females from 5 populations naturally infected by *Wolbachia* could discriminate between infected and uninfected males. Next, we performed experimental evolution under three selection regimes, corresponding to populations of spider mites that were either fully infected with *Wolbachia*, fully uninfected, or with intermediate infection frequency in males, to test if the evolution of pre-copulatory mating behaviour in response to CI is contingent upon the frequency of *Wolbachia* infection in the population.

## Materials and Methods

### Spider mite populations and rearing conditions

Seven *T. urticae* populations were used for these experiments: AMP, CH, COL, DC, DF, LOU, RF (Zélé et al. 2018a). All populations were collected in 2013 in Portugal, from different plants: AMP on *Datura* spp.; CH and RF on tomato (*Solanum lycopersicum*); COL and DF on bean (*Phaseolus vulgaris*); DC on zucchini (*Cucurbita pepo*) and LOU on eggplant (*Solanum melongena*). These populations were then established at the University of Lisbon, from 65 to 500 females, on bean plants (*Phaseolus vulgaris*, Fabaceae, var. *Enana*; Germisem Sementes Lda, Oliveira do Hospital, Portugal) under controlled conditions (25°C, photoperiod of 16L: 8D). The prevalence of *Wolbachia* was high (80 to 100%; (Zélé et al. 2018a) upon collection, but reached fixation in all populations and induced variable levels of CI in the laboratory (Zélé et al. *in prep.*).

### Experimental procedure

#### Mate choice in field-derived populations

5 naturally *Wolbachia*-infected *T. urticae* populations (AMP, CH, COL, DC, LOU) and their uninfected homologues were used to test if, within each population, uninfected females displayed a preference for uninfected or infected males. To create uninfected homologue populations, 30 adult females were placed in petri dishes containing bean leaf fragments on cotton wet with tetracycline solution (0.1 %, w/v) as described in (Zélé et al. in prep.). This treatment was applied continuously for three successive generations (Breeuwer 1997), followed by at least 20 generations of mass-rearing in an antibiotic-free environment, to prevent (or limit) potential side effects of antibiotic treatment (Ballard and Melvin 2007; Zeh et al. 2012). Before being used in all experiments, pools of 100 females were checked by PCR to confirm the *Wolbachia* infection status as described in (Zélé et al. 2018c).

*Wolbachia*-infected and -uninfected adult males and *Wolbachia*-uninfected quiescent females were separately isolated onto 8 cm^2^ leaf squares placed on water-saturated cotton from a subset of their base populations. The next day, quiescent females became virgin adults, roughly of the same age, while adult males had been isolated for *ca.* 24 hours, which guaranteed increased eagerness to mate (Krainacker and Carey 1990). Before the test, males of each population were painted with one of two distinct colours of water-based paint using a fine brush. An equal number of replicates per male type was assigned to each colour. The preference tests were done on 0.5 cm^2^ leaf discs (hereafter called “arenas”). Two males, from the same population but different infection status, were placed on each arena. The test started as soon as a *Wolbachia*-uninfected virgin female from the same population was added to the arena. Each preference test lasted for thirty minutes and the time until the beginning of mating-latency to copulation - and copulation duration were measured using a stopwatch (www.online-stopwatch.com). Simultaneously, the colour of the male that first copulated with the female was registered, and later assigned to a male type. The correspondence between male type and colour was only determined after observations to ensure observer blindness. Trials where no mating occurred for 30 minutes were excluded from the final analysis. In total, *ca.* 35 replicates per population were done (AMP: n=34; CH: n=33; COL: n=38; DC: n=32; LOU: n=37).

#### Establishment of populations for experimental evolution

Two subsets of each field-derived population (AMP, CH, COL, DC, DF, RF, LOU) were created by allowing the same founding individuals (n=100 females from each populations) to oviposit in two independent patches. One of these patches was treated with antibiotics to remove *Wolbachia* infection. This was done as previously described, except for the following details: 4 groups of 25 adult females per population subset were treated with tetracycline and mixed at the end of the treatment; the tetracycline-treated populations were maintained in absence of antibiotics for three generations before being used. As before, pools of 100 females from each population were checked by PCR to confirm the *Wolbachia* infection status (Zélé et al. 2018c) prior to the onset of experimental evolution. For one of the tetracycline-treated population (LOU), the PCR diagnostic for *Wolbachia* infection gave ambiguous results so we opted for excluding this population from subsequent experiments. One generation before starting experimental evolution, two base populations, infected or cured from *Wolbachia*, each replicated 5 times, were created by mixing 50 females from each of the 6 remaining *Wolbachia*-uninfected populations and from their 6 *Wolbachia*-infected homologues, respectively (i.e. each base population started with 300 females; see Electronic supplementary materials, Fig. S1). This procedure ensured a relatively high genetic diversity within all replicated populations, and a similar level of diversity across replicates.

#### Experimental Evolution

Each population of experimental evolution started by placing 200 females from each replicate of the base populations at 23.5°C in an experimental box (14×14×20cm) containing two bean plants (17 days old), whose stem was imbibed in wet cotton. A fresh bean plant was added to each experimental box after 7 days to avoid resource depletion. The eggs laid by the females in the experimental boxes hatched and reached adulthood within 14 days (i.e., generation time). At each generation, 200 young mated daughters were randomly picked from the old plants and transferred onto 2 fresh bean plants in a new experimental box. Three experimental regimes were created (Fig. 1): (a) a regime with *Wolbachia*-infected individuals only (hereafter called “iC” selection regime for “infected control”), (b) another with *Wolbachia*-uninfected individuals only (hereafter called “uC” selection regime for “uninfected control”) and (c) a regime consisting of *Wolbachia*-uninfected females and an even proportion of *Wolbachia*-infected and -uninfected males (hereafter called “uM” selection regime for “uninfected mixed”). In this latter regime, at each generation, 350 young quiescent females were randomly picked and placed on a bean leaf placed on water-saturated cotton, on which they emerged as adult virgins and were then let to choose between with 100 iC and/or 100 uM males for three days. 200 mated females were then transferred to fresh plants in a new box to build the next generation. For each selection regime, 5 independent replicates were maintained for 20 generations. Despite considerable care, one of the replicates of the mixed regime was contaminated by *Wolbachia*-infected females at generation 13. This replicate was thus excluded from the entire experiment. Consequently, only 4 replicates of all selection regimes were included in the experiment presented here. Data were obtained from tests performed at generations 12 to 15 of experimental evolution.

**Figure 1.**
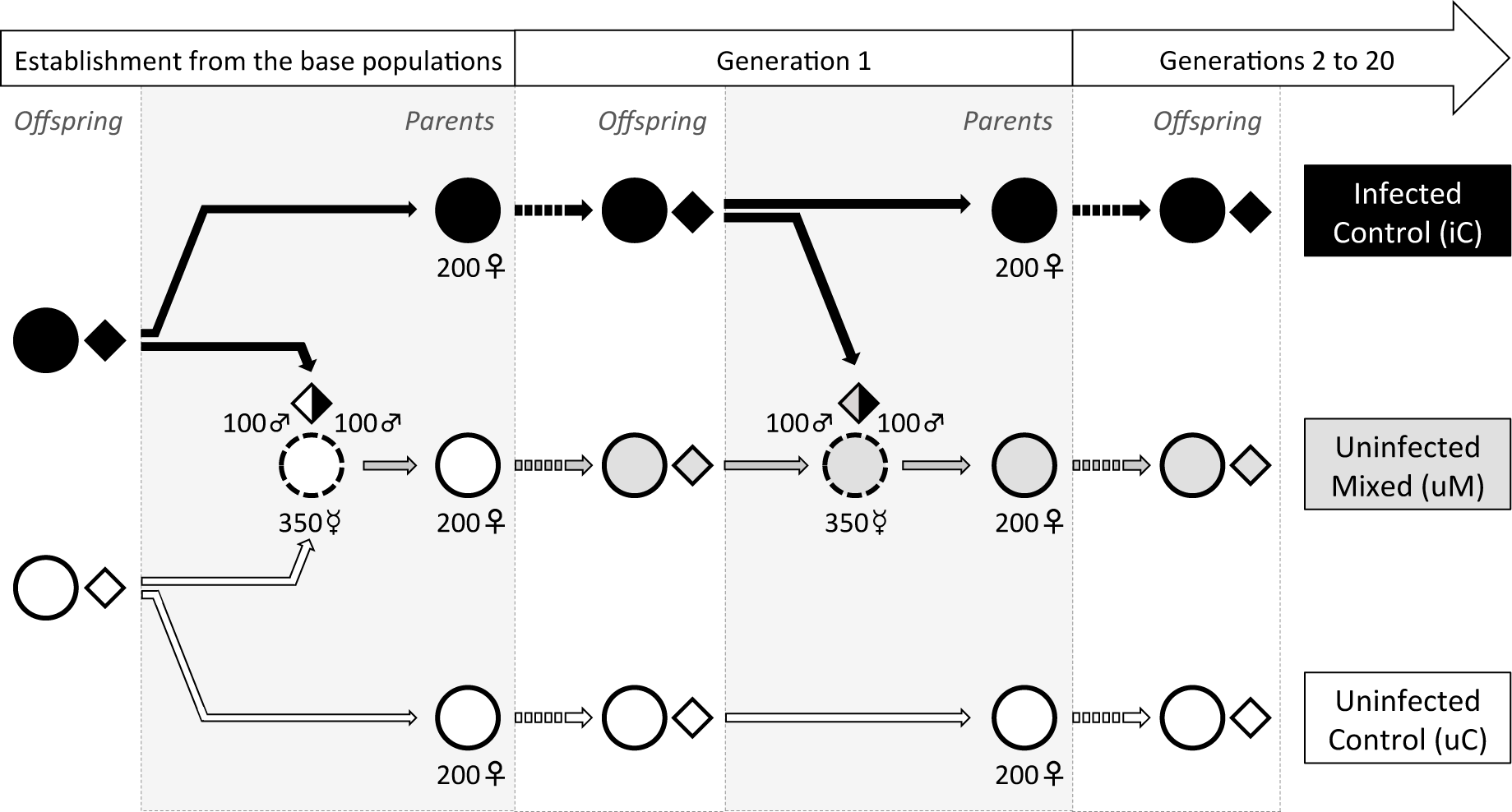
Procedure used for experimental evolution of spider mites under different infection scenarios. Shaded background: experimental manipulation of *Wolbachia* infection in males or in females used to create the next generation. The procedure in generations 2 to 20 is identical to that of generation 1. In the Uninfected-mixed regime, the infection status of males mating with uninfected females was controlled, while in all other treatments matings occurred before the transfers. Solid arrows: experimental transfers; White background and dashed arrows: offspring production and development. Circles: females; Diamonds: males; Solid-lined symbols: mated females (♀) and males (♂); Dashed-lined symbols: virgin females (☿); Black fill: *Wolbachia*-infected control (iC); Grey fill: *Wolbachia*-uninfected females mixed with -infected males (uM); White fill: uninfected control (uC). The entire procedure was repeated in 5 independent replicates.

#### Mate choice after experimental evolution

*Wolbachia*-uninfected females belonging to the control (uC) or mixed (uM) regimes, were given the choice between males of the uC and uM regimes, of the iC and uM regimes, or of the iC and uC regimes. To avoid an effect of preference due to differences in relatedness between and within replicates, females and males of each preference test belonged to different replicates: females from the replicates 1, 2, 3 and 4 mated with males from the replicates 2, 3, 4 and 1, respectively. The protocol followed here was similar to that of the first experiment except for two minor differences. First, males, like females, were isolated as quiescent from a subset of their base populations to ensure that all individuals were virgin and roughly of the same age. Second, trials where no mating occurred for 30 minutes were included in the final analysis, to test whether mating propensity (i.e., whether individuals mated during the time of the observations) evolved, as uninfected females could become less receptive to matings involving *Wolbachia*-infected males.

### Statistical Analyses

All analyses were carried out using the R statistical package (v. 3.0.3). Maximal models were simplified by sequentially eliminating non-significant terms (Crawley 2007), and the significance of the explanatory variables was established using chi-squared tests, in the case of discrete distributions, or F tests, in the case of continuous distributions (Bolker et al. 2008).

#### Mate choice in field-derived populations

Each population was analysed separately, since they were tested in a different time period, depending on spider mite availability and owing to excessive workload. To test for mate preference and an effect of colour on this choice in each population, we used Pearson’s Chi squared tests. Latency and duration of copulation were analysed using a cox proportional hazard mixed-effect model (coxme, package coxme), a non-parametric technique to analyse time-to-event data (e.g., time-to-death; Crawley 2007). In this analysis none of the data were censored, as non-mated females were excluded from the analysis (see above). Male type (infected or not with *Wolbachia*) was fit as a fixed explanatory variable, whereas day and colour of the chosen male were fit as random explanatory variables.

#### Mate choice after experimental evolution

To determine whether mating propensity and female mate choice were affected by the female selection regime (uM or uC), the type of preference test (choice between uM and uC, uM and iC or uC and iC), and/or their interaction, analyses were conducted using a generalized liner mixed-effect model (glmer, lme4 package) with a binomial error distribution. In both models, female selection regime and type of preference test were fit as fixed factors, whereas day, replicate nested within female selection regime, and colour assignment (i.e. the colour of each male type in an arena for the analysis of mating propensity, or the colour of the chosen male for the analysis of mate choice), were fit as random factors. To compare the outcome of preference tests to random mating, we performed a G-test of goodness-of-fit test (G.test, RVAideMemoire package).

To test for differences in mating latency and duration of copulation between selection regimes, the male chosen (uM, uC or iC) and the female selection regime (uM or uC) were fit as fixed explanatory variables, whereas type of preference test, day, replicate and colour of the chosen male were fit as random explanatory variables. Latency and duration of copulation were analysed as described above. Significant differences in factors with more than two levels were analysed using multiple comparisons with Bonferroni corrections (glht, package multicomp).

## Results

### Mating behaviour in field-derived populations

No effect of male colour on female choice was detected in any population (Table S1). Furthermore, uninfected females showed no significant preference for uninfected or infected males in any of the populations tested (AMP: X^2^_1_=0.12; P=0.73; CH: X^2^_1_=2.45; P=0.12; COL: X^2^_1_=0.42; P=0.52; DC: X^2^_1_=1.13; P=0.29; LOU: X^2^_1_=0.68; P=0.41; Fig. 2).

**Figure 2.**
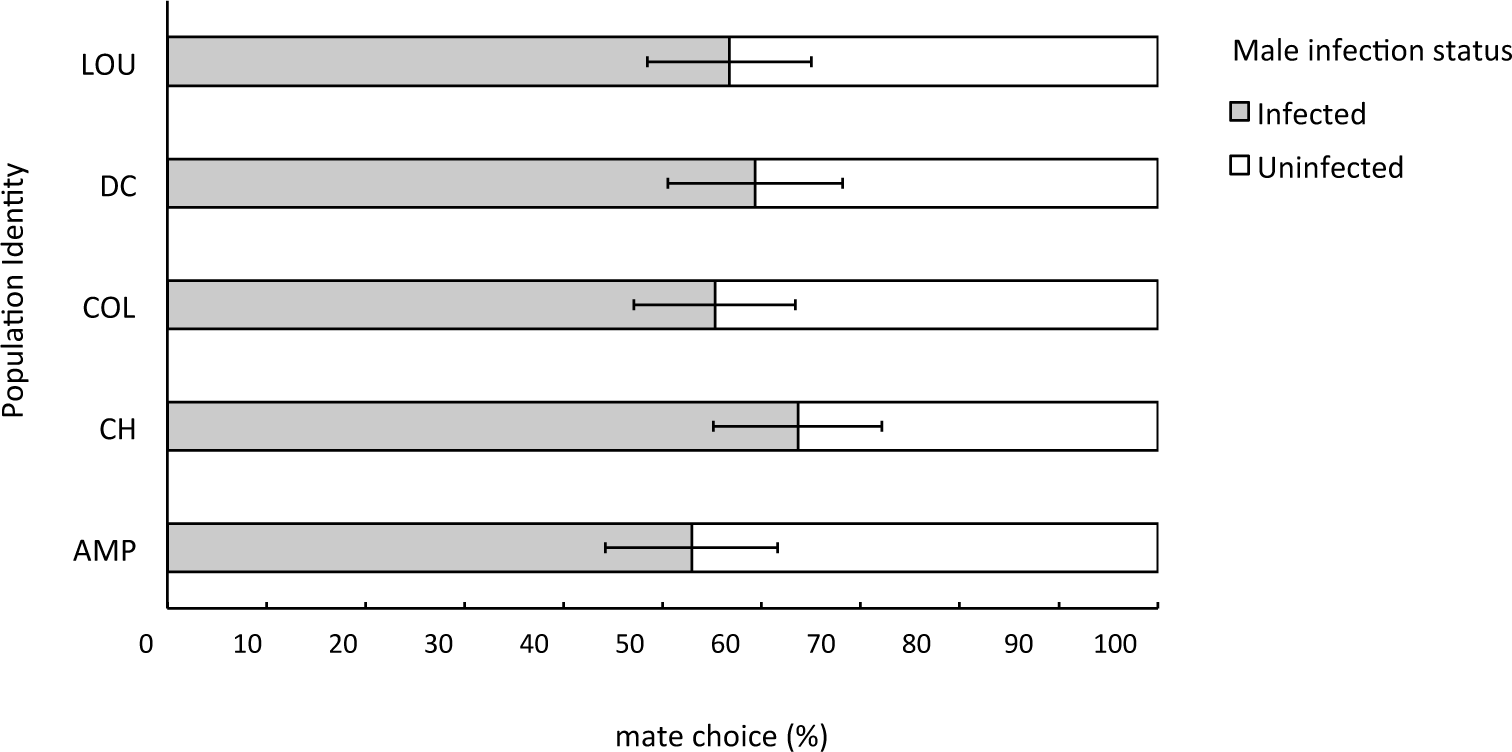
Mate choice in crosses between *Wolbachia*-uninfected females and *Wolbachia*-infected or -uninfected males in five populations of *T. urticae*. Bars represent means (± s.e.) percentage of *Wolbachia*-infected males (grey bars) or uninfected males (white bars) chosen by *Wolbachia*-uninfected females. Population identity: AMP, CH, COL, DC and LOU.

Moreover, latency to copulation with *Wolbachia*-infected or -uninfected males did not differ significantly (AMP: X^2^_1_=0.83; P=0.36; CH: X^2^_1_=3.21; P=0.07; COL: X^2^_1_=0.29; P=0.59; DC: X^2^_1_=0.34; P=0.56; LOU: X^2^_1_=0.005; P=0.95; Fig. 3a). Copulation duration did not differ significantly between crosses involving infected or uninfected males in AMP, CH, DC and LOU populations tested (AMP: X^2^_1_=0.16; P=0.69; CH: X^2^_1_=3.25; P=0.07; DC: X^2^_1_=0.02; P=0.89; LOU: X^2^_1_=0.21; P=0.65). However, in the COL population, copulations lasted longer with *Wolbachia*-infected males than with uninfected males (X^2^_1_=10.45; P=0.001; Fig. 3b).

**Figure 3.**
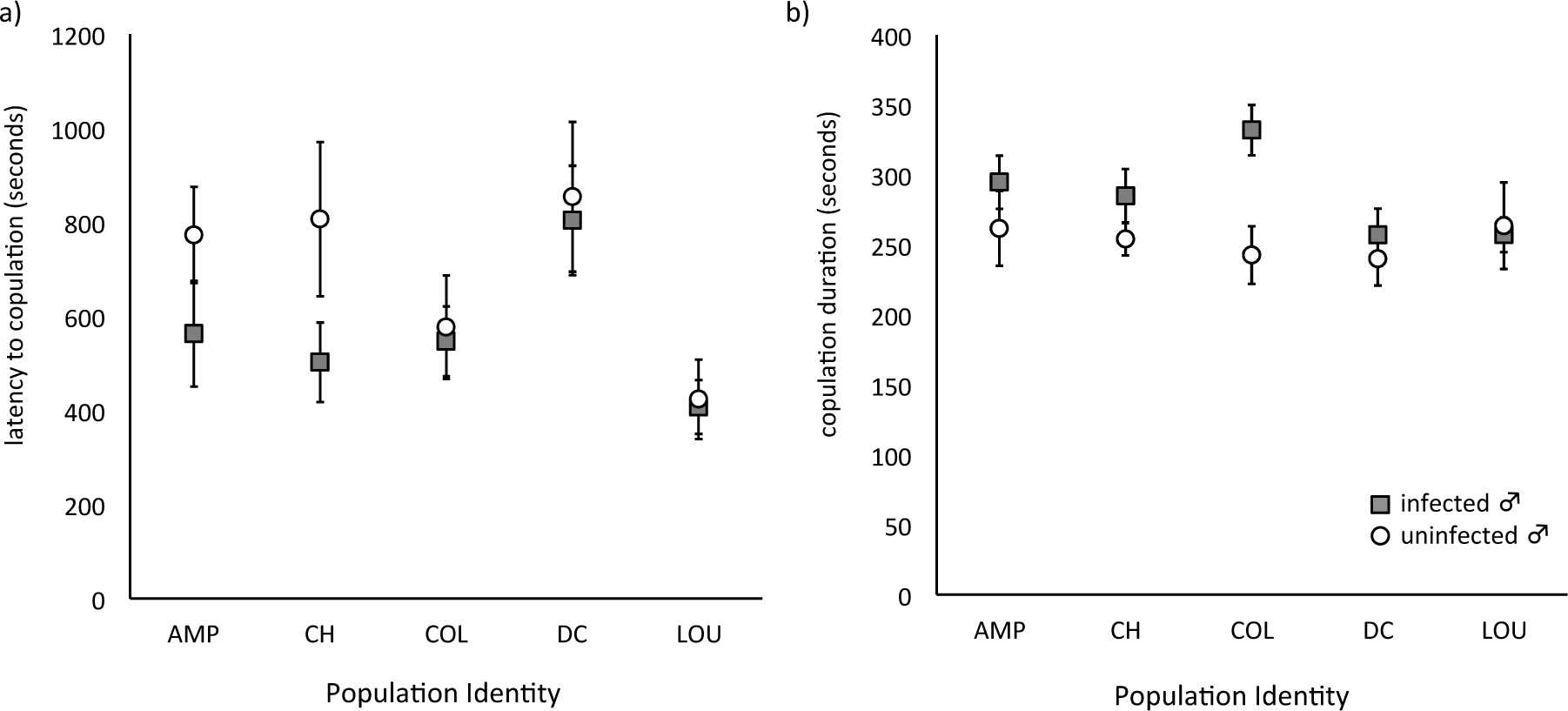
Latency to copulation (a) and copulation duration (b) of matings involving *Wolbachia*-uninfected females and *Wolbachia*-infected or -uninfected males of five populations of *T. urticae*. Mean (± s.e.) durations (in seconds) for *Wolbachia*-infected males (grey circles) and uninfected males (white squares). Population identity: AMP, CH, COL, DC and LOU.

### Mating behaviour after experimental evolution

The selection regime of the females tested, the type of preference test, and the interaction between these two factors did not significantly affect mating propensity (X^2^_1_=0.77, P=0.38; X^2^_2_=2.12, P=0.35 and X^2^_2_=1.03, P=0.60, respectively; Fig. 4a) nor mate choice (X^2^_1_=0.41, P=0.52; X^2^_2_=1.09, P=0.35 and X^2^_2_=1.01, P=0.60, respectively; Fig. 4b). Moreover, when comparing the preference tests to random mating using a goodness of fit test, no differences were found (Heterogeneity G: G=3.50, df=5, P=0.62; Pooled G: G_1_=2.77, df=1, P=0.10; Total G: G=6.27, df=6, P=0.39). Furthermore, no effect of female selection regime, of the type of male chosen, or of their interaction was found for latency to copulation (X^2^_1_=1.80, P=0.18; X^2^_2_=4.53, P=0.10 and X^2^_2_= 0.14, P=0.93, respectively; Fig. 5a). In contrast, copulation duration was significantly affected by the type of male chosen, but not by selection regime of the female, nor by the interaction between these factors (X^2^_2_=9.29, P=0.009; X^2^_1_=0.16, P=0.69 and X^2^_2_= 2.73, P=0.26, respectively; Fig. 5b). Indeed, females from all selection regimes engaged in longer matings with *Wolbachia*-infected males than with uninfected males from the control regime (uC vs iC: Z= −3.02, P=0.007), while no difference was found when comparing the other types of males (uC vs uM: Z=-1.96, P=0.12; iC vs uM: Z=1.18, P=0.47).

**Figure 4.**
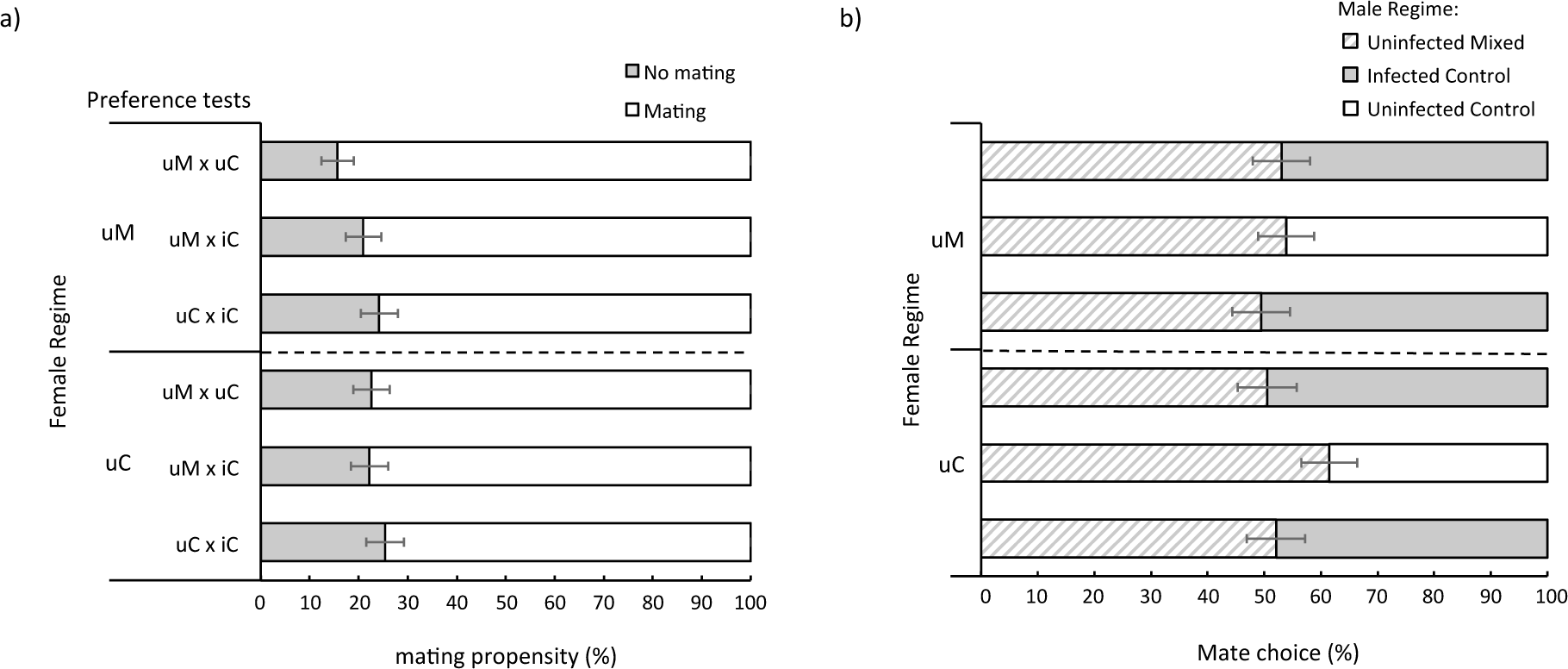
Mating propensity (a) and mate choice (b) of *Wolbachia*-uninfected females exposed to males from different selection regimes. *Wolbachia*-uninfected females from the control (uC) and the mixed regimes (uM) were given the choice between males from two different regimes (type of preference test): from uM, from uC or from iC (*Wolbachia*-infected control regime). In (a) bars represent mean (± s.e.) percentage of trials where mating occurred (white bars) or not (grey bars) within the time of the observation. In (b) bars represent mean (± s.e.) percentage of females choosing uC males (white bars), iC males (grey bars), or uM males (dashed bars).

**Figure 5.**
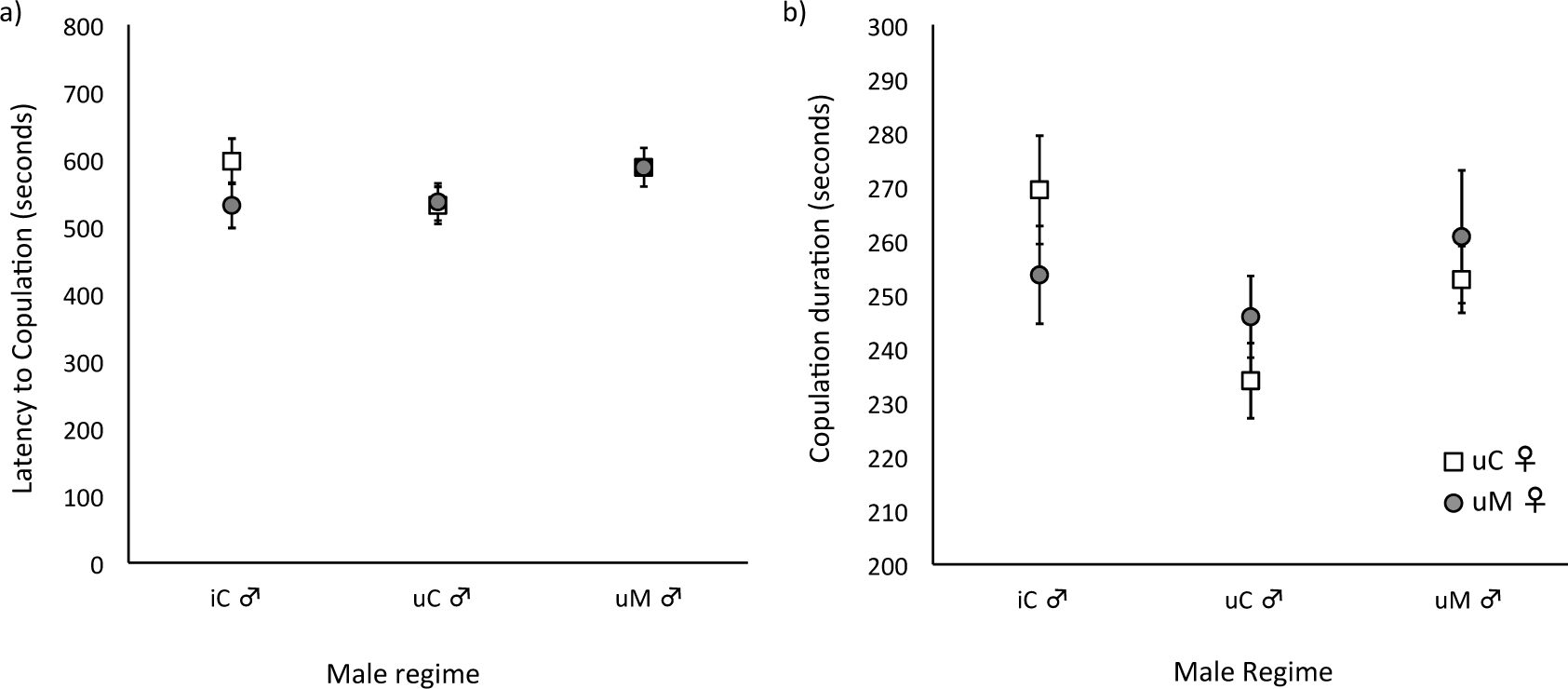
Latency to copulation (a) and duration of copulation (b) of matings between *Wolbachia*-uninfected females and males from different selection regimes. Mean (± s.e.) time (in seconds) for uC females (white squares) and uM females (grey circles). uM –*Wolbachia*-uninfected mixed selection regime; uC –*Wolbachia*-uninfected control regime; iC –*Wolbachia*-infected regime.

## Discussion

Here, we studied the mating behaviour of *Wolbachia*-uninfected females prior and after 12-15 generations of selection in environments with different frequencies of *Wolbachia* infection. Mate avoidance was not observed in any of the field-derived populations, nor did it evolve in the mixed-infection regime. Therefore, we found no evidence that spider mites collected from the field have pre-copulatory strategies to avoid *Wolbachia*-induced incompatibilities. Nevertheless, we found differences in copulation duration between infected and uninfected males in one out of 5 field populations, as well as between infected and uninfected males from the control regimes after experimental evolution. Conversely, we did not find that copulation duration differed between infected males from the control regime and uninfected males from the mixed-infection regime.

The higher duration of matings with *Wolbachia*-infected males, initially only present in the COL population, was recapitulated after experimental evolution: copulation duration was longer in matings with infected than with uninfected males from both control regimes. This suggests that infected males displaying longer copulation durations in the infected base populations increased in frequency during experimental evolution. Possibly, a longer copulation duration is advantageous for *Wolbachia*-infected males. Indeed, increased time of copulation has been implied in the insurance of paternity (Potter and Wrensch 1978; Satoh et al. 2001; Simmons 2001) and in an increase in the production of fertilized offspring in several species (Simmons 2001). Moreover, if infected males are able to fertilize more eggs than uninfected males when mated with uninfected females, this should result in an increased penetrance of CI. Accordingly, behavioural advantages conferred by *Wolbachia* to infected males, such as increased competitiveness and mating rate, has been shown in other species (Champion de Crespigny and Wedell 2006; Panteleev et al. 2007; but see Zhao et al. 2013). Alternatively, *Wolbachia*-infected males may mate longer to compensate for a decrease in sperm quality or quantity induced by *Wolbachia* (Snook et al. 2000; Champion de Crespigny and Wedell 2006; Lewis et al. 2011). In this case, we expect a decrease in the fertilized offspring of infected females in compatible crosses involving infected males, compared to those involving uninfected males due to an effect of *Wolbachia* on male fertility. However, no correlation between copulation duration and the number of female offspring (i.e., fertilized offspring in arrhenotokous spider mites) has been found in *T. urticae* (Satoh et al. 2001), suggesting that prolonged copulation may not be associated with increased fertility. Accordingly, the population with a higher copulation duration in matings with *Wolbachia*-infected males (COL) does not show significant differences in the sex-ratio of *Wolbachia*-infected and uninfected mites (Zélé et al*. in prep*.), which invalidates the first hypothesis. Moreover, infected males from the other field-derived populations (i.e. in which *Wolbachia* does not increase the copulation duration) do not lead to a more male-biased sex-ratio than uninfected males in compatible crosses (Zélé et al*. in prep*.), which suggests that *Wolbachia* does not lead to reduced sperm quality or quantity in our system, this way invalidating the second hypothesis. The benefits of such behaviour thus remain elusive and more studies are necessary to unveil the function of increased copulation duration in these circumstances.

In contrast to uninfected males from the control regime, uninfected males from the mixed-infection regime mated for as long as Wolbachia-infected males. This suggests that male competitive ability increased in the mixed infection regime, via selection on uninfected females. However, as no differences were observed between uninfected males from the control and from the mixed-infection regime, the evolved change in mating duration observed for males from the mixed regime is subtle. Still, in line with the results observed here, a study on reproductive interference between two spider mite species showed that *T. urticae* incompatible crosses with *T. evansi* did not elicit strong mate choice but heterospecific matings lasted less than conspecific ones (Clemente et al. 2016). Reproductive incompatibilities may thus generally result in changes in mating investment rather than in mating preference, which suggests that copulation duration is a more labile trait than mate choice in spider mites.

Indeed, no evidence for uninfected female mate choice between *Wolbachia*-infected and -uninfected males across field-derived populations was found here. This suggests that the ability to choose between males with different *Wolbachia* infection status is not common in *T. urticae* populations, and that the results obtained by Vala et al. (2004) are probably not representative of the reproductive behaviour of this species. Several factors may affect the probability that such choice evolves, such as the genotype, population structure and infection history of the host, as well as the *Wolbachia* strain (Engelstädter and Telschow 2009; Goodacre and Martin 2012). Possibly, these factors differ among studies, which may explain the differences found. Thus, to further understand the evolution of choice in host populations exposed to *Wolbachia*, one would need to compare this trait in populations that differ specifically in one of the above-mentioned factors. Here, we have controlled for the frequency of infection, maintaining it at an intermediate level to ensure a continuous selection pressure for choice. However, although avoidance of *Wolbachia*-infected males is expected to yield high benefits under these circumstances, *Wolbachia*-uninfected females evolving under this selection regime did not mate preferentially with *Wolbachia*-uninfected males. The lack of mate preference observed in the mixed regime could be explained by several, non-exclusive, mechanisms.

First, it might be due to an absence of cues necessary for the discrimination between *Wolbachia*-infected and uninfected males. Indeed, although, microbial infections (including with *Wolbachia*) have been shown to alter molecular cues used for mate recognition in diverse arthropod hosts (Beltran-Bech and Richard 2014; Engl and Kaltenpoth 2018), the capacity of many symbionts (including parasites) to avoid being detected by uninfected hosts or to manipulate the behaviour of infected host for their own benefit has also been thoroughly documented (Schmid-Hempel 2011). In many host species (reviewed by Zug and Hammerstein 2015), including *T. urticae* (Zhang et al. 2015), *Wolbachia* has evolved means to evade the host immune system. Likewise, it is likely that infected hosts are able to remain undetected by uninfected ones.

Another possibility is that pre-existing discrimination for a locally adapted trait, which is linked with the male infection status, is necessary for preference for compatible mates to evolve (Telschow et al. 2002, 2007). Thus, theoretically, uninfected females could evolve the ability to discriminate indirectly between *Wolbachia*-uninfected and -infected males via discrimination between related and unrelated males, respectively. Such scenario might be possible in *T. urticae* as mate discrimination based on relatedness has been shown in this species in absence of *Wolbachia* (Tien et al. 2011). Here, however, our experimental procedure was specifically designed to test for a direct effect of *Wolbachia*, hence not allowing for the expression of such indirect mechanism of preference (i.e. all individuals come from the same base populations and we always combined males and females from different replicate populations).

Alternatively, *Wolbachia*-induced cues to exert preference may be present in the population but females may not be able to perceive them, both before and after selection. Indeed, a preference allele could have been present in the field-derived populations but at a too low frequency, hampering its spread in the population. This is particularly expected if preference has a recessive genetic basis (Champion de Crespigny et al. 2005). Another possibility is that selection during experimental evolution might not have been strong enough for the allele to increase in frequency. Indeed, spider mites are haplodiploid (Helle and Sabelis 1985) and CI is incomplete in *T. urticae* (Gotoh et al. 2007; Zhang et al. 2016, Zélé et al. *in prep.*). Thus, females involved in incompatible crosses still pass on their genes via haploid sons and some females escape CI. Moreover, the introgression of the infected genotype in the uninfected population (i.e. iC males introgressed in uM selection regime) reduces the speed of evolution. Together, this is expected to result in a weaker selection pressure for the evolution of preference. Still, a theoretical model, with migration from a *Wolbachia*-infected mainland to (initially) uninfected island populations (i.e. a situation similar to that of our experimental design except that, in our study, only males migrate), predict that intermediate level of unidirectional CI can select for pre-mating isolation (Telschow et al. 2007). Moreover, spider mites have first male sperm precedence, i.e. only the first mating of a female is effective (Helle 1967). If this pattern cannot be disrupted, we expect the existence of a strong selection pressure on pre-copulatory strategies in this species.

Finally, mate choice may have not been observed because it trades-off with another beneficial trait or because its evolution is not a requisite for suffering reduced costs. For instance, if male quality is variable for other reasons, females may have to choose between mating with better quality or more compatible males (Colegrave et al. 2002; Neff and Pitcher 2005). The ability to avoid incompatible crosses could then be too costly to be maintained in an environment where incompatible crosses do not occur. This would explain our results, since the populations studied here were kept in the laboratory, fully infected, for *ca.* 24 generations before being tested for mate choice, and 30 more generations passed between these measurements and the observations done after experimental evolution. Another possibility is that the selective pressure applied here may also have led to the evolution of another trait that renders precopulatory mate choice unnecessary. For instance, spider mites may have evolved cryptic female choice or improved sperm competitive ability to avoid incompatible matings, as seen in other species (Price and Wedell 2008; Wedell 2013).

Our results show that assortative mating does not evolve in sympatry despite strong unidirectional post-zygotic barriers between populations. In the absence of complete post-copulatory isolation (e.g. incomplete unidirectional CI) gene flow between infected and uninfected individuals may prevent the evolution of reproductive isolation (Telschow et al. 2007). In this case, host speciation might be contingent upon the evolution of pre-copulatory strategies. The lack of evolved assortative mating here thus supports the hypothesis that maternally inherited symbionts that are able to manipulate the reproduction of their hosts do not necessarily lead to host speciation. Moreover, our results suggest that hosts may not evolve behavioural traits that function as a strong barrier to the spread of *Wolbachia* in host populations, such as mate choice. Hence, the maintenance of infection polymorphisms in natural populations may rather hinge upon host abundance, population structure and migrations (Hancock et al. 2011; reviewed in Engelstädter and Telschow 2009), as well as factors known to affect *Wolbachia* transmission, fitness effects and/or CI levels, such as environmental variables (e.g. temperature, resource availability and/or quality; (Corbin et al. 2017; Zélé et al. 2018b; Zhu et al. 2018) and hosts traits (e.g. genetic background, development time, aging; Merçot and Charlat 2004; Yamada et al. 2007; Sun et al. 2016; LePage et al. 2017).

Finally, such finding also has important implication, as mate preference should not hamper the success of deliberate introductions of *Wolbachia* into mosquito populations in several regions worldwide with the potential to control vector-borne disease agents (Hoffmann et al. 2011, 2014; Nguyen et al. 2015).

## Author contributions

Experimental conception and design: LRR, FZ, SM; acquisition of data: LRR, FZ, IS; statistical analyses: LRR; paper writing: LRR, FZ, SM, with input from all authors. All authors have read and approved the final version of the manuscript.

## Acknowledgments

We are grateful to Diogo Godinho and João Alpedrinha for useful discussions and comments and all Mite Squad members for unconditional support in the laboratory and elsewhere. This work was funded by an FCT-ANR project (FCT-ANR//BIA-EVF/0013/2012) and an ERC consolidating grant (COMPCON GA 725419) to SM. FZ was funded through FCT Post-Doc fellowship (SFRH/BPD/125020/2016). The authors declare no conflict of interest.

